# NAD modulates DNA methylation and cell differentiation

**DOI:** 10.1101/2020.07.31.231555

**Authors:** Simone Ummarino, Mahmoud A. Bassal, Yanzhou Zhang, Andy Joe Seelam, Ikei S. Kobayashi, Marta Borchiellini, Alexander K. Ebralidze, Bon Q. Trinh, Susumu S. Kobayashi, Annalisa Di Ruscio

## Abstract

Nutritional intake impacts the human epigenome by directing epigenetic pathways in normal cell development via as yet unknown molecular mechanisms. Consequently, imbalance in the nutritional intake is able to dysregulate the epigenetic profile and drive cells towards malignant transformation. Herein, we present a novel epigenetic effect of the essential nutrient, NAD. We demonstrate that impairment of DNMT1 enzymatic activity by NAD-promoted ADP-ribosylation, leads to demethylation and transcriptional activation of *CEBPA* gene, suggesting the existence of an unknown NAD-controlled region within the locus. In addition to the molecular events, NAD treated cells exhibit significant morphological and phenotypical changes that correspond to myeloid differentiation.

Collectively, these results delineate a novel role for NAD in cell differentiation and indicate novel nutri-epigenetic strategy to regulate and control gene expression in human cells.

## Introduction

Malnutrition and obesity are associated to epigenetic dysregulation thereby promoting cellular transformation and cancer initiation (Avgerinos, Spyrou et al., 2019, Birks, Peeters et al., 2012). A prolonged exposure to high-fat diet, poor nutrition and insults from environmental toxicants, all contribute to the epigenetic transgenerational inheritance of the obesity (King & Skinner, 2020). The degree of obesity, in term of body weight, is a well-documented risk factor for hematopoietic disease and cancer (Strom, Yamamura et al., 2009, Tedesco, Qualtieri et al., 2011). Together, these evidences highlight the importance of balanced micronutrient intake in order to preserve cell specific epigenetic programming and prevent anomalies that can potentially result in malignant transformation (Montgomery & Srinivasan, 2019, Yilmaz, Atilla et al., 2020).

In the last decade, numerous studies focusing on establishing a link between nutrition and epigenetics, leaded to the concept of “Precision Nutrition”; a translational approach based on the use of dietary compounds to direct epigenetic changes and drive normal cellular development (Zeisel, 2020). Natural compounds, like vitamins C and D, have been shown to slow pathological processes through their impact on the epigenome (Bunce, Brown et al., 1997, Nur, Rath et al., 2020). Similarly, nutri-epigenomic approaches have been shown to prevent several disease conditions including cancer (Di Tano, Raucci et al., 2020, Meroni, Longo et al., 2020). Nevertheless, the molecular mechanisms by which nutrients modulate the epigenome of healthy or cancer cells is largely unknown.

Nicotinamide adenine dinucleotide (NAD) is a dietary compound essential for life, and a coenzyme implicated in cellular redox reactions (Rajman, Chwalek et al., 2018). Maintenance of adequate levels of NAD is critical for cellular function and genomic stability (Ralto, Rhee et al., 2020). Few reports have shown that NAD precursors such as vitamin B3 (or nicotinic acid, NA) and nicotinamide (Nam) are able to drive cell differentiation in leukemic cell lines (Ida, Ogata et al., 2009, Iwata, Ogata et al., 2003), and impair cell growth. However, the molecular mechanism participating in these morphological changes remain unknown.

DNA methylation is a key epigenetic signature involved in transcriptional regulation, normal cellular development, and function (Jones, 2012). Methyl groups are added to the carbon 5 of cytosines in the contest of CpG dinucleotides by specialized enzymes the DNA methyltransferase enzymes (DNMT1, 3A and 3B). While the bulk of the genome is methylated at 70–80% of its CpGs, CpG islands (CGI), that are clusters of CpG dinucleotides generally proximal to the transcription start sites (TSSs) of most human protein-coding genes, are mostly unmethylated in somatic cells. Numerous studies have established a link between aberrant promoter DNA methylation and gene silencing in diseases such as cancer (Herman & Baylin, 2003, Jones & Baylin, 2002).

NAD is also the substrate of Poly-(ADP) Ribose Polymerase 1 (PARP1) a nuclear protein that plays a pivotal role in gene regulation, and chromatin remodeling (Hageman & Stierum, 2001, Ray Chaudhuri & Nussenzweig, 2017). PARP1 utilizes NAD as a source of ADP-ribose moieties to assemble ADP-ribose polymers (PAR) and coordinate epigenetic modifications including DNA methylation (Ciccarone, Zampieri et al., 2017, Reale, Matteis et al., 2005). Several experimental data support a PARP1-mediated inhibition of DNA methyltransferase 1 (DNMT1) activity in human cell lines (Fang, Bi et al., 2015, Witcher & Emerson, 2009). These findings suggest a role for NAD in altering and, or facilitating modulation of DNA methylation, even if a direct link between demethylation and NAD treatment has not been established (Ciccarone, Valentini et al., 2014, Di Ruscio, Ebralidze et al., 2013).

Herein, we present a novel function of NAD, the ability to specifically demethylate and induce the expression of the hematopoietic master regulator, CCAAT/enhancer binding protein alpha (*CEBPA*) gene locus. The demethylation effect correlates with a total and local increase of ADP-ribose polymers (PAR) at the *CEBPA* promoter, thus supporting a NAD/PARP1/DNMT1 axis in which local inhibition of DNMT1, results in site-specific demethylation and transcriptional activation.

Our findings indicate NAD as a novel epigenetic modulator that counteracts the widespread epigenetic reprogramming concurring to obesity and cancers, and provide the first nutritional-based therapy for clinical interventions in these conditions.

## Materials and Methods

### Cells and Cell Culture

K562 cell line was purchased from ATCC and grown in RPMI medium supplemented with 10% fetal bovine serum (FBS), in the absence of antibiotics at 37°C, 5% CO_2_. The K562-*CEBPA*-ER line was grown in 12 well plate in phenol red–free RPMI 1640 (ThermoFisher Scientific, Cat. No. 11835030), supplemented with 10% Charcoal stripped FBS (Sigma Aldrich, Cat. No. F6765), and 1 µg/mL puromycin, beginning at a density of 0.2 × 10^6 cells/mL. 1µM estradiol (Sigma Aldrich, Cat. No. E2758) was added from a 5-mM stock solution in 100% ethanol to induce *CEBPA*-ER nuclear translocation and a corresponding amount of ethanol (0.02%) to mock-treated cells as controls. Viable cells excluding trypan blue were enumerated every day and used for the experiment (D’Alo, Johansen et al., 2003, Umek, Friedman et al., 1991).

### NAD treatment

K562 cells were incubated with 0.1, 0.5, 1, 1.5 or 10 mM of NAD (Sigma Aldrich) or vehicle (milliQ water) for four days at 37°C. Cells were counted every day and cell pellets were collected to perform all the downstream analysis.*Colorimetric NAD assay*. The BioVision NAD/NADH Quantification Colorimertic Kit was used according to the manufacturer’s instructions (BioVsion). Briefly, K562 cells were homogenized by two cycles of freezing and thawing in 400μl of BioVsion NAD/NADH extraction buffer. The homogenate was filtered using BioVision 10-kD cut-off filters (10000 g, 25 min, 4°C). To detect only NADH content, NAD was decomposed by heating 200µl of the homogenate. The homogenate of decomposed and non-decomposed samples was distributed in a 96 well plate, the developer solutions was added to the samples. The absorbance (OD 450nm) was acquired for 30 minutes using the VICTOR Multilabel Plate Reader (Perkin Elmer).

### K562 Wright Giemsa staining

Approximately 2 ×10^4^ per each sample, were spotted on a slide using the cytospin at 400 rpm for 5 min. The cells were then stained with the Wright Giemsa solutions kit (CAMCO STAIN PAK, pc#702) according to manufacturer’s instructions.

### Nitroblue Tetrazolium (NBT) assay

Nitroblue blue tetrazolium (NBT) analysis was performed using 5×10^5^ cells incubated in a 1 mL solution containing phosphate-buffered saline (PBS), NBT (Sigma), and 0.33 µM phorbol myristate acetate (PMA) for 20 minutes at 37°C. The reaction was then stopped by incubation on ice. Cells were immediately fixed on slides by cytocentrifugation and counterstained with 0.5% safranin O in 20% ethanol.

### Immunofluorescence

Cells were fixed with PFA 2% (Paraformaldehyde/MeOH), washed with 1X Phosphate-buffered saline (PBS), and permeabilized with 0.5 % Triton X100. After blocking with 7% Goat-serum, for 30 min, cells were incubated with primary antibody Anti poly (ADP-ribose) polymer (1:400 Abcam, ab 14459) overnight at 4°C, covered from the light. The following day, cells were washed with 1X PBS, and incubated with secondary antibody goat anti-mouse (1:500, Alexa Fluor 555) for 1hr, re-washed, and nuclei counterstained with Prolong gold antifade mountant already containing DAPI (Thermo Fisher Scientific). Samples were analysed on a Leica DM 5500B Microscope with a 100W high-pressure mercury lamp. Images were assembled and contrast-enhanced using Image J as per manufacturer’s recommendations.

### RNA isolation and qRT-PCR analyses

Total RNA isolation was carried out using TRIzol (Thermo-Fisher Scientific), as previously described (23). All RNA samples used in this study were treated with DNase I (10 U of DNase I per 3 µg of total RNA; 37 °C for 1 hr; in the presence of RNase inhibitor). After DNase I treatment, RNA samples were extracted with acidic phenol (Sigma, pH 4.3) to eliminate any remaining traces of DNA. Taqman based qRT-PCR was performed using the one step Affymetrix HotStart-IT qRT-PCR Master Mix Kit (Affymetrix USB) and 50 ng of total RNA per reaction. Amplification conditions were 50°C (10 min), 95°C (2 min), followed by 40 cycles of 95°C (15s) and 60°C (1 min). Target gene amplification was calculated using the formula 2^^−ΔΔCt^ as described (23), primer and probe sequences are listed in supplementary Tab 1.

### DNA isolation

Cell pellets, resuspended in a homemade lysis buffer (0.5% SDS, 25 mM EDTA pH 8, 10 mM TRIS pH 8, 200 mM NaCl), were initially treated with RNase A (Roche) for 20 minutes at 37°C and then Proteinase K (Roche) overnight at 65°C. High quality genomic DNA was extracted by Phenol:chloroform:isoamyl Alcohol 25:24:1, pH:8 (Sigma) and precipitated with Isopropanol the following day. Genomic DNA was resuspended in Tris 1mM, EDTA 10mM (TE) pH 8 and stored at 4°C.

### Western blotting analysis

Whole-cell lysates from approximately 2 ×10^5^ cells per each sample were separated on 15% SDS-PAGE gels and transferred to a nitrocellulose membrane. Immunoblots were all blocked with 5% nonfat dry milk in Tris-buffered saline, 0.1% (TBS-T) prior to incubation with primary antibodies. The Anti-poly (ADP-ribose) polymer (1:1000 Abcam, ab14459) was stained overnight at 4°C. For PARP1 and DNMT1 protein analyses, equivalent amount of whole-cell lysates were separated on 7 % SDS-PAGE gels and transferred to a nitrocellulose membrane. Immunoblots were stained overnight with the following primary antibodies: Anti-PARP1 (1:1000 Active motif, 39559), Anti-DNMT1 (1:1000, Abcam, ab19905). All secondary horseradish peroxidase (HRP)-conjugated antibodies were diluted 1:5000 and incubated for 1hr at room temperature with TBST/ 5% milk. Immuno-reactive proteins were detected using the Pierce^®^ ECL system (Thermo Scientific #32106).

### Bisulfite Sequencing and Analysis

DNA methylation profile of *CEBPA* locus was analyzed by bisulphite sequencing as previously described (Di Ruscio et al., 2013). Briefly, high molecular weight genomic DNA (1µg) was subjected to bisulfite conversion using the EZ DNA Methylation-Direct kit (Zymo Research) following the manufacturer’s instructions. Polymerase chain reactions (PCR) on bisulfite converted DNA was performed with FastStart Taq DNA Polymerase (Roche) in the following conditions: 95°C (6 min) followed by 35 cycles at 95°C (30 s) 53-57°C (1 min) 72°C (1 min), and a final step at 72°C (7 min). Primers and PCR conditions for bisulfite sequencing are summarized in supplementary Tab 2. After gel purification, cloning into PGEM T-easy vector (Promega) and transformation in E. coli Competent Cells JM109 (Promega), 9 positive clones analyzed by Sanger sequencing for each sample. Only clones with a conversion efficiency of at least 99.6% were considered for further processed by QUMA: a quantification tool for methylation analysis (http://quma.cdb.riken.jp/) (Kumaki, Oda et al., 2008).

### Chromatin immunoprecipitation

ChIP was performed as previously described (Zhang, Alberich-Jorda et al., 2013). Briefly, K562 cells were crosslinked with 1% formaldehyde (formaldehyde solution, freshly made: 50 mM HEPES-KOH; 100 mM NaCl; 1 mM EDTA; 0.5 mM EGTA; 11% formaldehyde) for 10 min at room temperature (RT) and 1/10^th^ volume of 2.66 M Glycine was then added to stop the reaction. Cell pellets were washed twice with ice-cold 1X PBS (freshly supplemented with 1 mM PMSF). Pellets of 2 ×10^6^ cells were used for immunoprecipitation and lysed for 10 minutes on ice and chromatin fragmented using a Bioruptor Standard (30 cycles, 30 sec on, 60 sec off, high power). Each ChIP was performed with 10µg of antibody, incubated overnight at 4°C. A slurry of protein A or G magnetic beads (NEB) was used to capture enriched chromatin, which was then washed before reverse-crosslinking and proteinase K digestion at 65°C. Beads were then removed in the magnetic field and RNase treatment (5µg/µl Epicentre MRNA092) performed for 30 minutes at 37°C. ChIP DNA was extracted with Phenol:chloroform:isoamyl Alcohol 25:24:1, pH:8 (Sigma) and then precipitated with equal volume of isopropanol in presence of glycogen. DNA pellet was dissolved in 30µl of TE buffer for following qPCR analyses. The following antibodies were used for ChIP: Anti-DNMT1 (Abcam, ab19905), Anti-poly (ADP-ribose) polymer (Abcam, ab14459), normal mouse IgG (Millipore 12-371b) and normal rabbit IgG (Cell Signaling 2729S). Fold enrichment was calculated using the formula 2 (-ΔΔCt (ChIP/non-immune serum)). Primer sets used for ChIP are listed in supplementary Table 3.

### Immunostaining for FACS analysis

Anti-CD15-APC (Thermal Fisher Scientific, Cat. No. 17-0158-42), anti-CD14-FITC (Thermal Fisher Scientific, Cat. No. 11-0149-42) and anti-CD11b-Pacific blue (BioLegend, Cat. No. 101224) were incubated with 1×10^6^ K562 cells (vehicle or NAD treated) at 1:100 ratio. Cells were pre-incubated with anti-Fc receptor antibody (Thermal Fisher Scientific, Cat. No. 14-9161-73) at 1:20 ratio to block Fc receptor before staining. Zombie red staining (BioLegend, Cat. No. 423109) was used as cell viability dye during FACS analysis. Cells were fixed using 2% PFA (Sigma, Cat. No. 158127) before performing FACS analysis. Cell acquisition and analysis were performed on BD LSRFortessa (BD biosciences, CA, USA) using BD FACSDiva™ software (BD Bioscience). Analysis was performed using Flowjo software (Flowjo LLC, USA).

### Annexin V staining

FITC Annexin V Apoptosis Detection Kit I (BD Bioscience) was used to determine the percentage of K562 undergoing apoptosis upon NAD treatment. All samples were prepared following the manufacturer’s instructions. Briefly, cells were collected every day, washed twice with cold PBS and then resuspended in 1x Binding buffed at a concentration of 1×10^6^ cells/ml. Cells were incubated with 5µl fluorescein isothiocyanate (FITC) annexin V and 5µl Propidium Iodide for 15 min at room temperature in darkness. Analyses of cells viability and apoptosis were performed on BD LSR Fortessa (BD biosciences, CA, USA) using BD FACSDiva™ software (BD Bioscience). The data analysis was performed using Flowjo software (Flowjo LLC, USA).

### Seahorse analysis

Mito Stress Test (Agilent Seahorse, 103015-100) assay was run as per manufacturers’ recommendations. Briefly, on the day of assay, counted and PBS washed cells were suspended in XF Assay media (Agilent Seahorse Bioscience) pH adjusted to 7.4 ± 0.1 supplemented with 4.5 g/L glucose (Sigma-Aldrich G7528), 0.11 g/L sodium pyruvate (Sigma-Aldrich) and 8 mM L-glutamine (Sigma-Aldrich). 1×10^5^ cells were added to each well of XFe24 Cell-Tak (Corning) pre-coated culture plates and then slowly centrifuged for incubation at 37°C in a non-CO_2_ incubator. Oxygen consumption rate was measured at baseline using a Seahorse XFe24 according to standard protocols and after the addition of oligomycin (100 μM), carbonyl cyanide-4-(trifluoromethoxy) phenylhydrazone (FCCP, 100 μM) and rotenone and antimycin A (50 μM). Fold change was determined by normalizing raw values to the average of the second basal reading.

### Statistical analysis

All bisulfite sequenced clones were analyzed by Fisher’s exact test. For mRNA qRT-PCR, *p-*values were calculated by t-test in GraphPad Prism Software. For both the analysis, values of *p* <0.05 were considered statistically significant (\**p* < 0.05; \*\**p* < 0.01; \*\*\**p* < 0.001). The Mean ± SD of duplicates is reported.

## Results

### NAD inhibits cancer cell growth in a dose-dependent manner and drives accumulation of intracellular poly ADP-ribose polymers

NAD precursors drive myeloid differentiation and impair cell growth (Ida et al., 2009, Iwata et al., 2003). To examine whether similar effects could be mediated by NAD, K562 cells were cultured following a single addition of NAD or vehicle to the media, and tracked over four days (**Fig. 1a**). Cells were counted every day and cell pellets collected for downstream analyses (**Fig. 1a, b**). Inhibition of the cell growth, was observed across all the tested NAD concentrations in a dose-dependent manner, with the strongest effect at 10 mM, 96 hours upon treatment (**Fig. 1b**). Notably, this inhibition was not associated with apoptosis as demonstrated by the Annexin V staining, showing high viability (≈ 85%) of NAD-treated cells *versus* untreated (**Fig. S1a**). Consistently, the NAD/NADH content in the 10mM NAD treated cells, displayed nearly eightfold increase as compared to the baseline, already 24 hours after treatment (**Fig. 1c**). Provided that NAD is partially, utilized as a source of ADP-ribose units by PARP1 to build linear and branched poly ADP-ribose (PAR) polymers, NAD-treated and untreated K562 were stained with an anti-PAR antibody and examined by immunofluorescence to monitor the accumulation of PAR. Consistently, 24 hours upon NAD treatment, cells displayed an intense fluorescence signal in treated as compared to untreated cells, owing to the increased PAR synthesis and accumulation (**Fig. 1d**).These results mirrored the effects induced by 10-minutes treatment with hydrogen peroxide (H_2_O_2_), a known DNA damaging agent (Blenn, Althaus et al., 2006, Ryabokon, Cieslar-Pobuda et al., 2009, Valdor, Schreiber et al., 2008), associated with PAR production and therefore used as a positive control (**Fig. S1b**). Overall, these findings supported a PAR accumulation driven by NAD. As a further validation, PAR levels were analyzed by western blot. The strongest PAR band was detected on the first day and gradually decreased in the following days (**Fig. 1e**) likely due to PARs degradation by poly (ADP-ribose) glycohydrolases (PARGs) or similar pathway-related enzymes (O’Sullivan, Tedim Ferreira et al., 2019).

**Figure 1.**
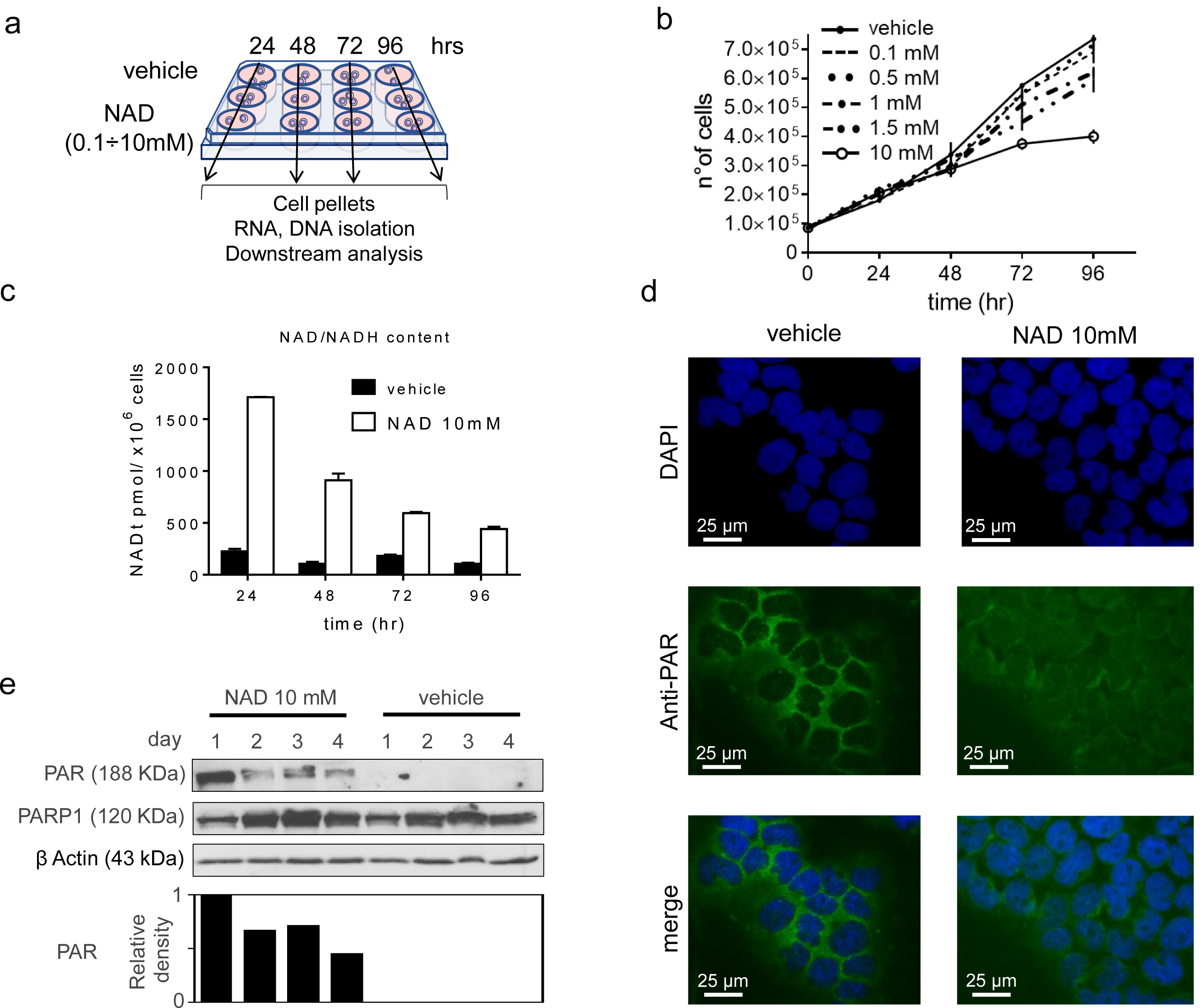
NAD inhibits cancer cell growth in a dose-dependent manner and drives accumulation of intracellular poly ADP-ribose polymers. (**A**) Schematic of the experiment. K562 cells were cultured at different concentration of NAD:0.1, 0.5, 1, 1.5, 10 mM or vehicle. Cell pellets, RNA and DNA samples were collected at different time point, 24, 48, 72, 96 hrs. (**B**) K562 growth curves in presence of NAD or vehicle. Cells were counted every 24 hrs for four days (**C**) The NAD/NADH content measured by colorimetric assay. The absorbance was measured at 450 nm every 24 hrs from the addition of NAD (10mM) to the cell culture media. NAD ratio was calculated according to the manufacturer’s instructions (BioVision). (**D**) Immunofluorescence of PARs in K562 supplemented with of NAD (10mM) or vehicle after 24 hrs. (**E**) PAR and PARP1 protein levels in K562 cells treated with NAD. The immunoblot band densities is measured using ImageJ and normalized by β-Actin.

Collectively these data demonstrate that NAD inhibit cell growth and mediates accumulation of intracellular PAR as early as 24 hours upon treatment.

### NAD treatment induces CEBPA distal promoter demethylation

A PARP1-mediated inhibition of DNMT1 activity in human cell lines has been reported (Fang et al., 2015, Reale et al., 2005). Therefore, we reasoned that increase of NAD, a substrate of PARP1, could modulate genomic methylation. To this end, we investigated the methylation dynamics of the well-studied methylation-sensitive gene *CEBPA* in K562 cells, following treatment with 10mM NAD (Hackanson, Bennett et al., 2008, Zhang et al., 2013) *CEBPA* is a master transcription factor in the hematopoietic system, the loss or inhibition of which can result in block of differentiation and granulopoiesis, contributing to leukemic transformation. *CEBPA* promoter is aberrantly methylated in ∼30% and ∼51% of patients with chronic myeloid leukemia and acute myeloid leukemia, respectively (Hackanson et al., 2008, Iwata et al., 2003, Tenen, 2003). As *CEBPA* promoter, encompassing the −1.4 kb to −0.5 kb regions from the transcriptional start site (TSS), is methylated in K562, we decided to assess the impact of NAD treatment on DNA methylation profile. Using bisulfite sequencing, we surveyed three distinct regions located at −0.8 kb (−557; −857), −1.1kb (− 895; −1.122), −1.4 kb (−1.120; −1.473) upstream to the TSS of *CEBPA* (**Fig. 2a**). NAD treatment led to concomitant decrease of DNA methylation levels within the distal promoter region (−0.8 kb) (**Figs. 2b, c and S2a**) which equaled 44% reduction at 48 hours and dropped to 60%, 72 hours after NAD addition. These levels bounced back to a mild 17% decrease after 96 hours suggesting a dynamic re-establishment of DNA methylation levels within the site (**Fig. 2b, c**). In agreement with our earlier findings (**Fig. 1d,e**), wherein the strongest accumulation of PARs was observed 24 hours post-NAD treatment (**Fig. 1d,e**), these results seem to indicate that the additional 24 hours were required to inhibit DNMT1 enzymatic activity and promote the methylation changes observed over the 48 and 72 hour time points (**Fig. 2b,c**). Unexpectedly, only minor changes in the distal promoter I (−1.1kb) and II (−1.4kb) were detected at 72 hours, suggesting a certain specific modality of NAD-mediated demethylation (**Figs. 2d, e and S2a**). Consistent with previous reports, DNA methylation within the −1.1kb and −1.4kb regions, does not correlate with *CEBPA* expression in both K562 and AML samples using conventional hypomethylating drugs (Hackanson et al., 2008).

**Figure 2.**
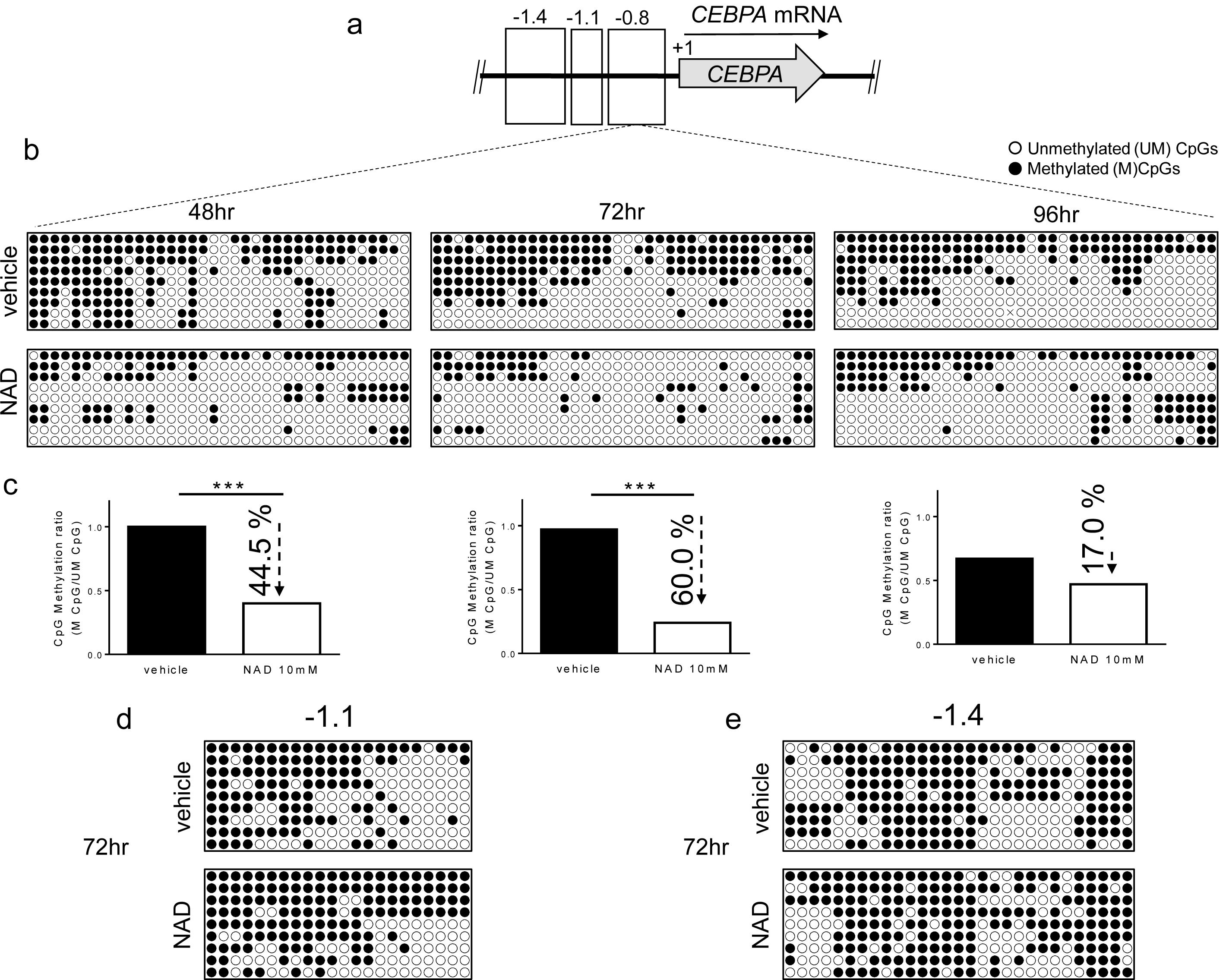
DNA methylation patterns of *CEBPA* upon NAD (10mM) or vehicle treatment. (**A**) Schematic representation of *CEBPA* locus. The three regions analysed in the promoter of *CEBPA* located at −0.8 kb (−557; −857), −1.1 kb (−895; −1.122) or −1.4 kb (−1.120; −1.473) from the TSS (+1) of the gene. (**B, C**) The methylation status of the distal promoter (the −0.8 kb region) was analysed at the three indicated time points. 9 clones were analysed, and lollipop graphs were generated using QUMA software. CpG methylation ratio consisting in methylated CpGs divided by unmethylated CpGs, was calculated by QUMA software. (**D, E**) Methylation status of distal promoter I (−1.1 kb) and distal promoter II (−1.4 kb) analysed 72hrs upon NAD (10mM) addition. Lollipop graphs were generated as described. (n=9 clones). All bisulfite sequenced clones were analysed by Fisher’s exact test, *:*p*<0.05; **:*p*<0.01; **:*p*<0.001

Taken together these data demonstrate that the NAD-induced *CEBPA* promoter demethylation relies on a PAR-dependent mechanism which impairs DNMT1 activity

### NAD treatment enhances CEBPA mRNA transcription in K562 by a PARP1-dependent mechanism

DNA methylation is a key epigenetic signature involved in gene regulation. To investigate whether NAD-induced demethylation of *CEBPA* distal promotor was associated with increased levels of *CEBPA* transcriptional activation, we measured the *CEBPA* expression by qRT PCR in cells treated with 10 mM NAD (**Fig. 3a**), over multiple time points. Upregulation of *CEBPA*, 72- and 96-hour-upon treatment was observed only in cells treated with the highest NAD concentration (**Figs. 3a and S2b**). These results parallel *CEBPA* upregulation at 72 and 96 hours following demethylation of the distal promoter using the standard hypomethylating agent 5-aza-2’-deoxycytidine in K562 cells (Hackanson et al., 2008). As the only region sensitive to NAD-induced demethylation effect corresponded to *CEBPA* distal promoter, while nearly no changes occurred in the two upstream regions (−1.4 kb and −1.1kb) we reasoned the involvement of epigenetic regulators to account for this site selectivity. Previous studies have reported that PARP1 assembled ADP-ribose polymers are able to impair DNMT1 activity in human and murine cell lines (Reale et al., 2005). In following these findings, we hypothesized a mechanism wherein the NAD-induced production of PAR would specifically inhibit DNMT1 activity at *CEBPA* distal promoter, without affecting the more upstream regions. To test this hypothesis, we firstly verified the levels of PARP1 and DNMT1 were not influenced by NAD at both expression and protein levels (**Figs. 3b and S2c**). Secondly, we performed quantitative Chromatin Immunoprecipitation (ChIP) with anti-PAR and anti-DNMT1 antibodies, 24 hours upon NAD treatment (**Fig. 3c-e**), given the strongest increase of PAR polymers at that specific time point (**Fig. 1d, e**). As expected, the *CEBPA* distal promoter region exhibited over 1.6-fold enrichment of PAR polymers than the vehicle treated cells, unlike the distal promoter II (**Fig. 3d, e**) in which the polymers were absent. Interestingly, DNMT1 distribution between the distal promoter and the regions more upstream was unchanged (**Fig. 3e**), suggesting the same accessibility of DNMT1 for both sites, and a potential impairment of the enzymatic activity at the distal promoter due to the presence of the PAR polymers.

**Figure 3.**
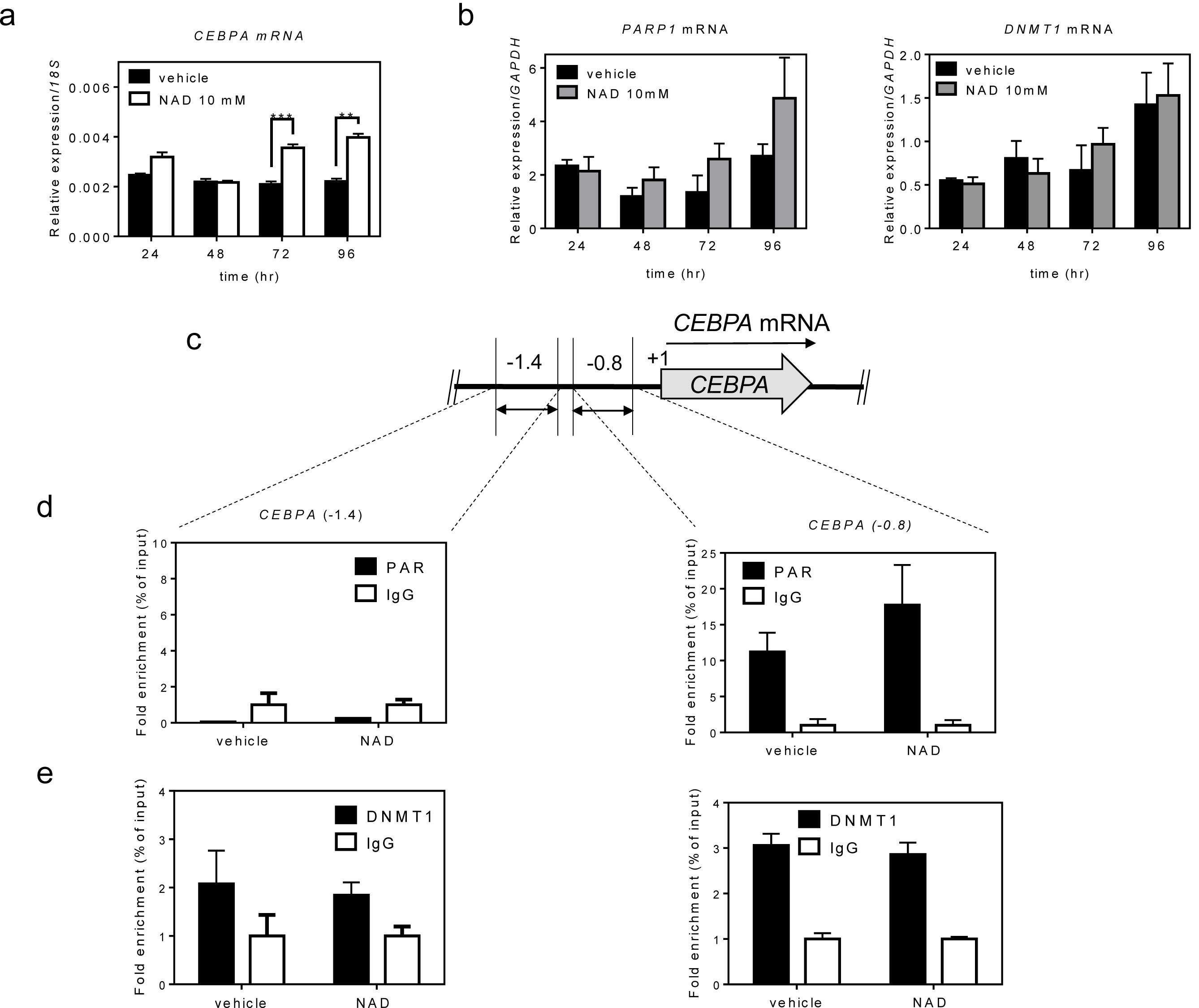
NAD treatment enhances *CEBPA* transcription in K562 by a PARP1-dependent mechanism. Panel (**A**) shows *CEBPA mRNA* levels upon 4 days treatment with NAD. The graph represents the average of two independent experiments (n=2). Panel (B) shows *PARP1* and *DNMT1 mRNA* levels upon 4 days treatment with NAD. Chromatin was collected to perform ChIP assays with antibodies to PAR, DNMT1 and IgG (**C-E**). (**C**) Schematic of the *CEBPA* promoter regions screened by ChIP-qPCR analysis respectively at −1.4 kb and −0.8 kb from the TSS (double-headed arrows). (**D**) ChIP using PAR antibody and qPCR analysis of regions −1.4 kb (left panel) and −0.8 kb (right panel). (**E**) ChIP using DNMT1 antibody and qPCR analysis of regions −1.4 kb (left panel) and −0.8 kb (right panel). Error bars indicate ± S.D. *:*p*<0.05; **:*p*<0.01; **:*p*<0.001

Collectively, these results indicate a PARP1-dependent demethylating mechanism boosted by NAD levels and enabling inhibition of DNMT1 activity in selected loci.

### NAD induces myeloid differentiation

As previously reported NAD-precursors such as NA and other niacin-related compounds induce differentiation in immortalized cell lines, such as K562 and HL60 (Ida et al., 2009, Iwata et al., 2003). These findings prompted us to assess morphological changes upon NAD treatment. Wright Giemsa staining of K562 treated with 10mM NAD or vehicle revealed striking morphological changes four days after treatment (**Fig. 4a**). Specifically, vehicle treated cells exhibited a homogeneous population of round-shaped cells, with round or oval cell nuclei, whereas NAD-treated cells were more heterogeneous, with a higher cytoplasm:nucleus ratio, eccentrically located reniform nuclei with dense regions of heterochromatin and numerous vacuoles resembling a monocytic-macrophagic morphology. Additionally, NAD treatment leads to increases in nitroblue tetrazolium (NBT)-positive cells and expression of CD11b and CD14 surface markers indicating that NAD promotes monocytic-macrophagic differentiation in K562, whilst the absence of CD15 expression ruled out a shift toward the granulocytic lineage (**Fig. 4b, c**) (Federzoni, Humbert et al., 2014). Hence, despite the reactivation of *CEBPA* mRNA, which is a master regulator of granulocytic differentiation, the expected morphological changes were not detected in NAD treated cells, although we could confirm increased expression of both CD15 and CD11b and not CD14 upon ectopic expression of CEBPA protein as already shown previously (Federzoni et al., 2014, Perrotti, Cesi et al., 2002) (**Fig. S3a**). These results are not surprisingly since the oncogenic fusion protein: BCR-ABL, that is constitutively expressed in K562, suppresses *CEBPA* translation thus leading to transcriptional suppression of the granulocyte colony-stimulating factor receptor G-CSF-R and other myeloid precursor cells critical for granulocytic differentiation (Perrotti et al., 2002). Along with these data, we confirmed the absence of *CEBPA* protein by western blot analysis on K562 NAD-treated cells (data not shown).

**Figure 4.**
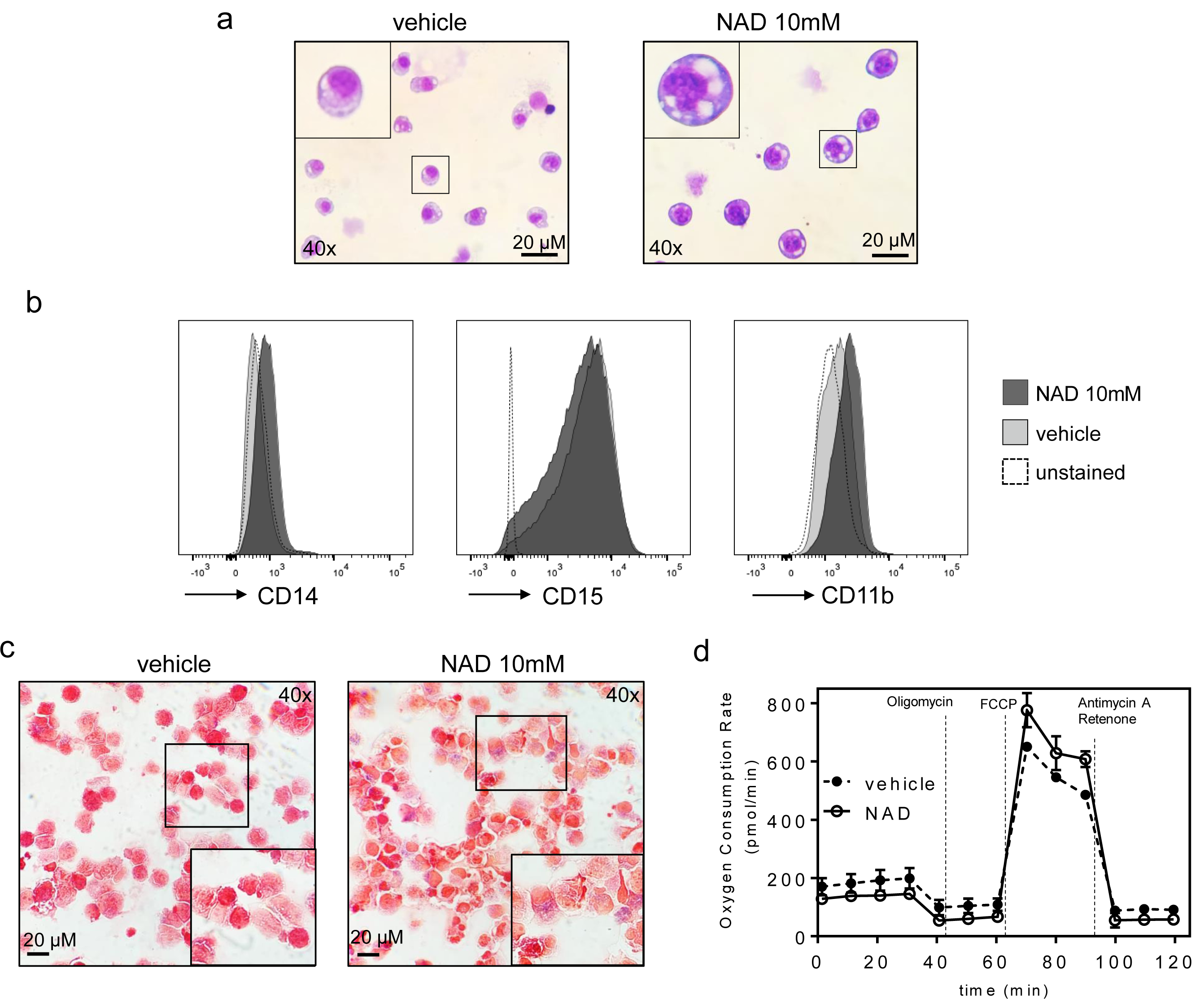
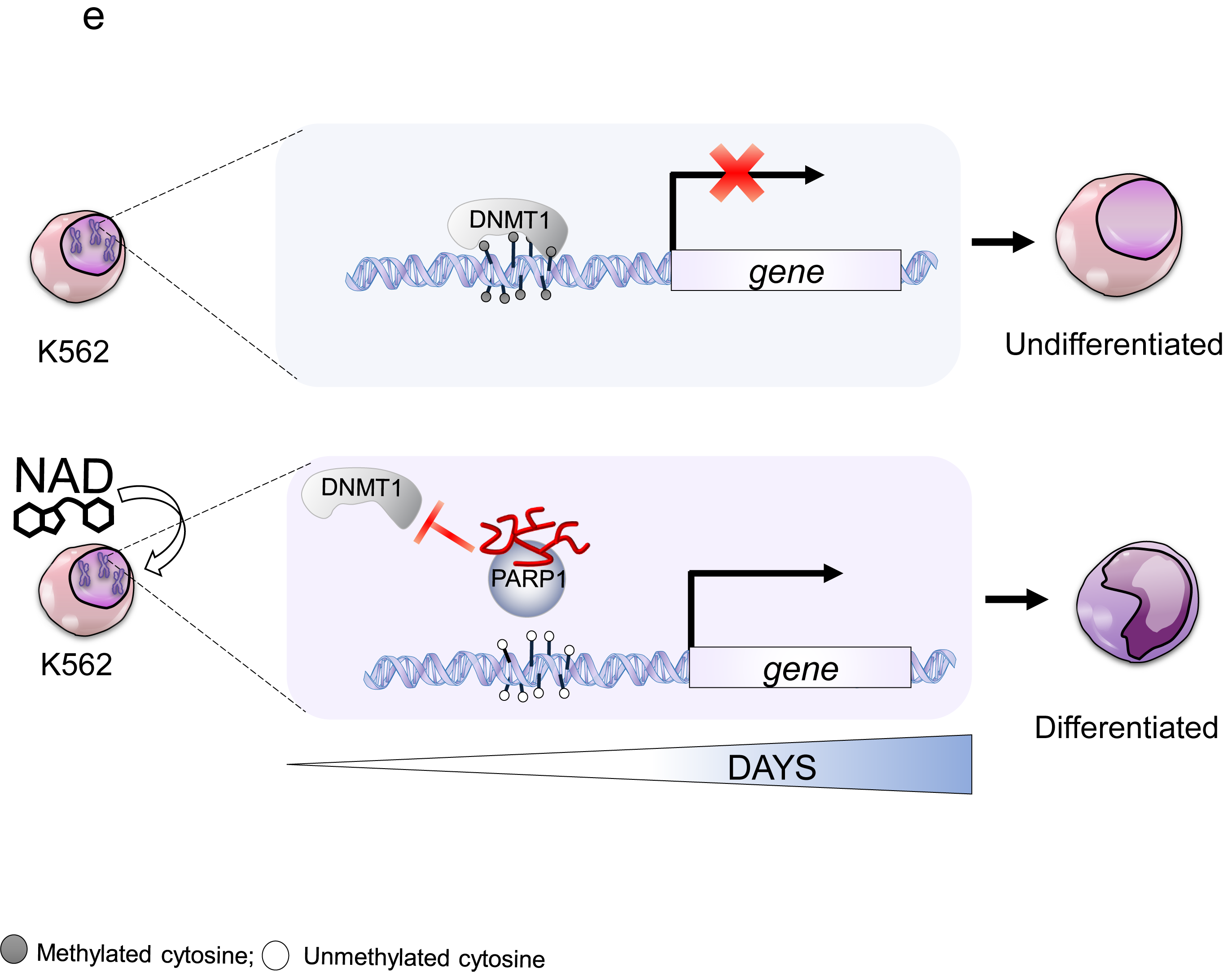
NAD induces myeloid differentiation in K562. (**A**) Wright Giemsa staining showing morphological changes between NAD-treated and control cells after four days. (**B**) Increase in the surface markers CD15, CD14 and CD11b upon NAD treatment (**C**) NBT positive staining detected by small blue dots after counterstaining the cells with safranin. A magnification is shown in the rectangle. (**D**) Seahorse XF analysis of K562 mitochondrial stress response in cells treated with NAD or vehicle. The figure represents the mean of two biological replicates (n=2). Error bars indicate ± S.D. (**E**) Model showing the molecular mechanism of *CEBPA* gene reactivation by NAD. *CEBPA* is epigenetically silenced in K562. DNMT1 ensure a constant methylated status of *CEBPA* promoter (upper part). The NAD supplementation to K562 cell culture, boosts PARP1 to produce ADP-ribose polymers leading to DNMT1 inhibition (bottom part). The ultimate effect is *CEPBA* re-activated transcription.

### NAD treatment improves mitochondrial OXPHOS function

NAD has been previously demonstrated to restore mitochondrial function in aged mice and increase the intracellular ratio of NAD^+^/NADH, a critical cellular balance required for the Sirtuin 1 (SIRT1) mediated activation of mitochondrial biogenesis (Chalkiadaki & Guarente, 2015, Khan, Auranen et al., 2014). To further investigate the NAD contribution to the mitochondrial function of K562, the Mito Stress Test was performed using a Seahorse XFe24 (**Fig. 4d**). Basal oxygen consumption rate (OCR) is used as a surrogate measure of mitochondrial function since mitochondria utilize oxygen to generate mitochondrial ATP. Our results show that NAD-treated K562 cells displayed a marginal increase in maximal oxygen consumption in response to Carbonyl cyanide-4 (trifluoromethoxy) phenylhydrazone (FCCP) stress. This translated to a 1.3-fold improvement in normalized maximal reserve capacity after only four days of co-incubation with NAD. Albeit a marginal change in maximal reserve capacity post NAD-treatment was observed, these results still highlight the significance NAD treatment plays on improving mitochondrial health and perhaps contributing to the changes described. The entire profile of K562 NAD-treated does not depart drastically from K562 untreated, but the increment of ORC emerging after the injection of FCCP, indicate variations in respiration capacity of K562 NAD-treated *versus* untreated, subjected to the same mitochondrial stimuli.

## Discussion

This study explores the demethylation impact brought about by NAD treatment. On the example of the *CEBPA* gene locus, silenced by DNA methylation in the leukemia model used herein, we carried out a molecular and biological dissection of the potential mechanism implicated in NAD-induced demethylation. We demonstrate that impairment of DNMT1 enzymatic activity, as a result from NAD-promoted ADP-ribosylation, leads to loss of *CEBPA* promoter methylation and corresponding transcriptional activation of *CEBPA* mRNA thereby revealing an unknown NAD-controlled region within the *CEBPA* locus.

NAD is regarded as a potential antiaging molecule, the levels of which tend to decline over our lifetime, yet the molecular mechanisms linking low NAD levels to aging are only partially understood (Bonkowski & Sinclair, 2016, Lautrup, Sinclair et al., 2019). As a critical substrate of SIRT and PARP enzyme family members, that are involved in multiple epigenetic pathways such as acetylation, ADP-ribosylation and DNA methylation, fluctuations of NAD levels may alter chromatin remodeling (Bai, 2015, Chalkiadaki & Guarente, 2015). An additional epigenetic role for NAD, independently of its partnering enzymes, has also been hypothesized by few reports wherein age- or nutrition-related decline of NAD levels were associated with the acquisition of abnormal DNA methylation profiles at specific loci (Chang, Zhang et al., 2010, Kane & Sinclair, 2019). *In vitro* evidence have also shown that ADP-ribosyl polymer impair DNMT1 enzymatic activity (Reale et al., 2005) and an ADP-ribosylation transcriptional control for the *P16* and *TET1* genes has been demonstrated (Ciccarone et al., 2014, Witcher & Emerson, 2009). To date over 2300 proteins, including DNMT1, have been reported as ADP-ribosylated (http://ADPriboDB.leunglab.org) but how ADP-ribosylation preserves the unmethylated state of certain regulatory sequences, remains elusive (Vivelo, Wat et al., 2017). In every instance studied, we demonstrate that NAD treatment induces production of PAR polymers, site-specific demethylation of *CEBPA* distal promoter and results in transcriptional activation of *CEBPA* mRNA in K562 cells (**Figs. 2,3**). These results led to hypothesize a site-selective demethylation mechanism wherein the NAD-induced production of PAR polymers inhibits DNMT1 activity at *CEBPA* distal promoter by preventing DNMT1 interaction with the CGI, as described in the depicted model (**Fig. 4e**). The co-occurrence of PARs and DNMT1 on the distal promoter, but not on the distal promoter II, suggests a PAR-mediated specific inhibition of DNMT1 and reveals a NAD-responsive element on *CEBPA* promoter (**Fig. 3**). Intriguingly, the morphological changes along with the pronounced NBT staining and the positive shift of CD11b and CD14 surface markers, in addition to the improved mitochondrial function, seems to point to a monocytic-macrophagic-like transcriptional activation program initiated by NAD treatment (**Fig. 4**).

In conclusion, this study bridges a nutritional intervention to a molecular observation: increase of NAD levels in a cancer cell line results in local correction of DNA methylation. These data, therefore, provides a nutritional-guided approach for the prevention and the clinical management of cancers or other conditions associated with alteration of DNA methylation, potentially linked to decreased NAD levels.

## Acknowledgements

This work was supported by the National Institution of Health R00 CA188595, the Italian Association for Cancer Research (AIRC) Start-up Grant N.15347, the Giovanni Armenise-Harvard Foundation Career Development Award to ADR; the National Institution of Health (R01CA169259 and R01CA240257) and Harvard Stem Cell Institute Blood Program (DP-0110-12-00) to SSK; the NCI K01CA222707 was awarded to BQT.

## Author contributions

ADR supervised the project. ADR and SU conceived and designed the study and wrote the manuscript; SU, MAB, YZ, AJ, ISK, MB, SSK, performed experiments; BQT, MAB, SSK and AKE provided valuable suggestions about the project, and BQT, MAB, SSK critically reviewed the manuscript.

## Data and materials availability

All data and materials are available in the main text or the supplementary materials.

---

**Supplementary figure 1.**
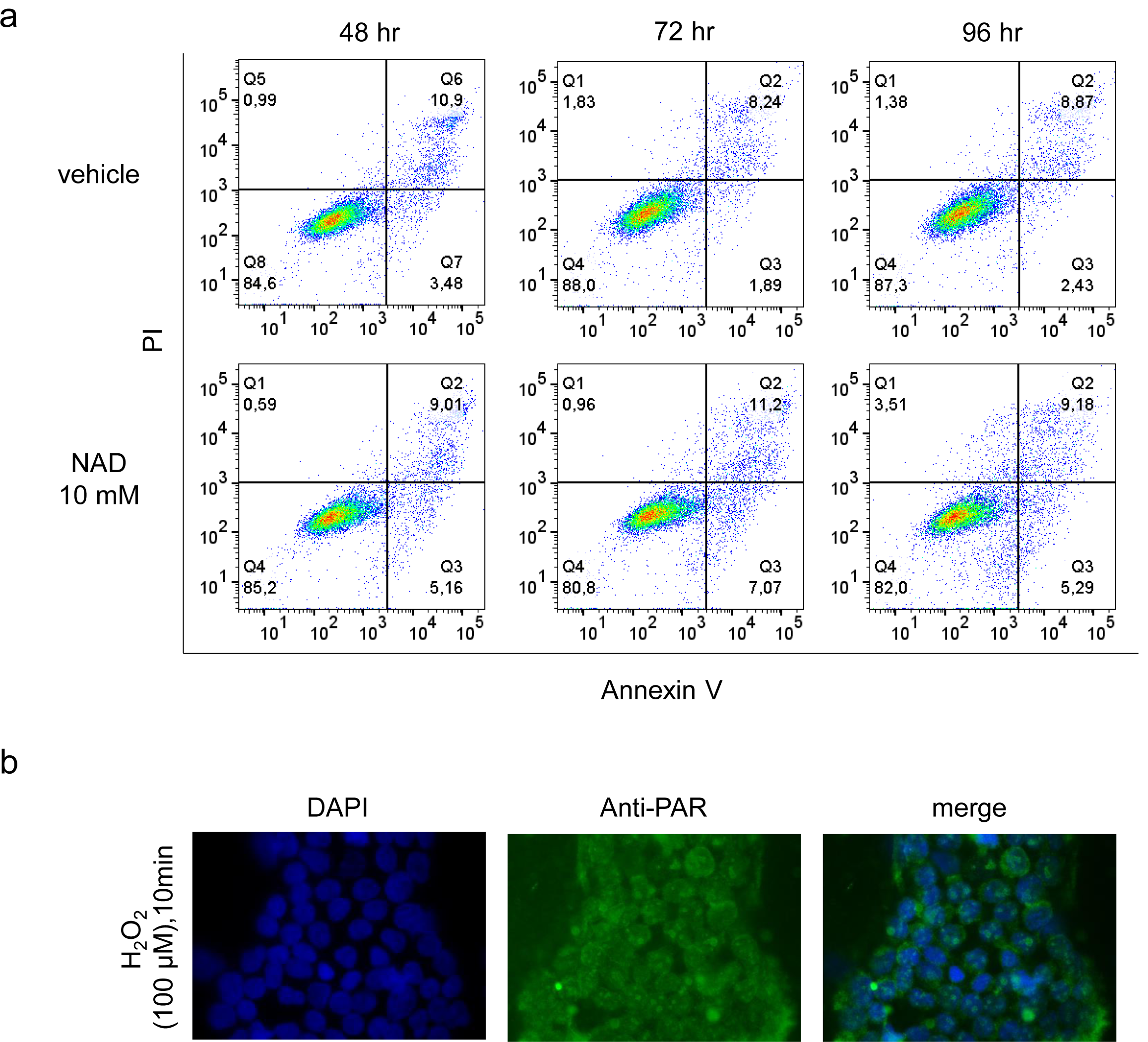
Cell viability upon NAD treatment. (**A**) K562 cells viability and apoptosis analyses upon NAD or vehicle (water) treatment. (**B**) Immunofluorescence analysis of PARs formation induced by 10 min-treatment with H_2_O_2_ (100µM) in K562.

**Supplementary figure 2.**
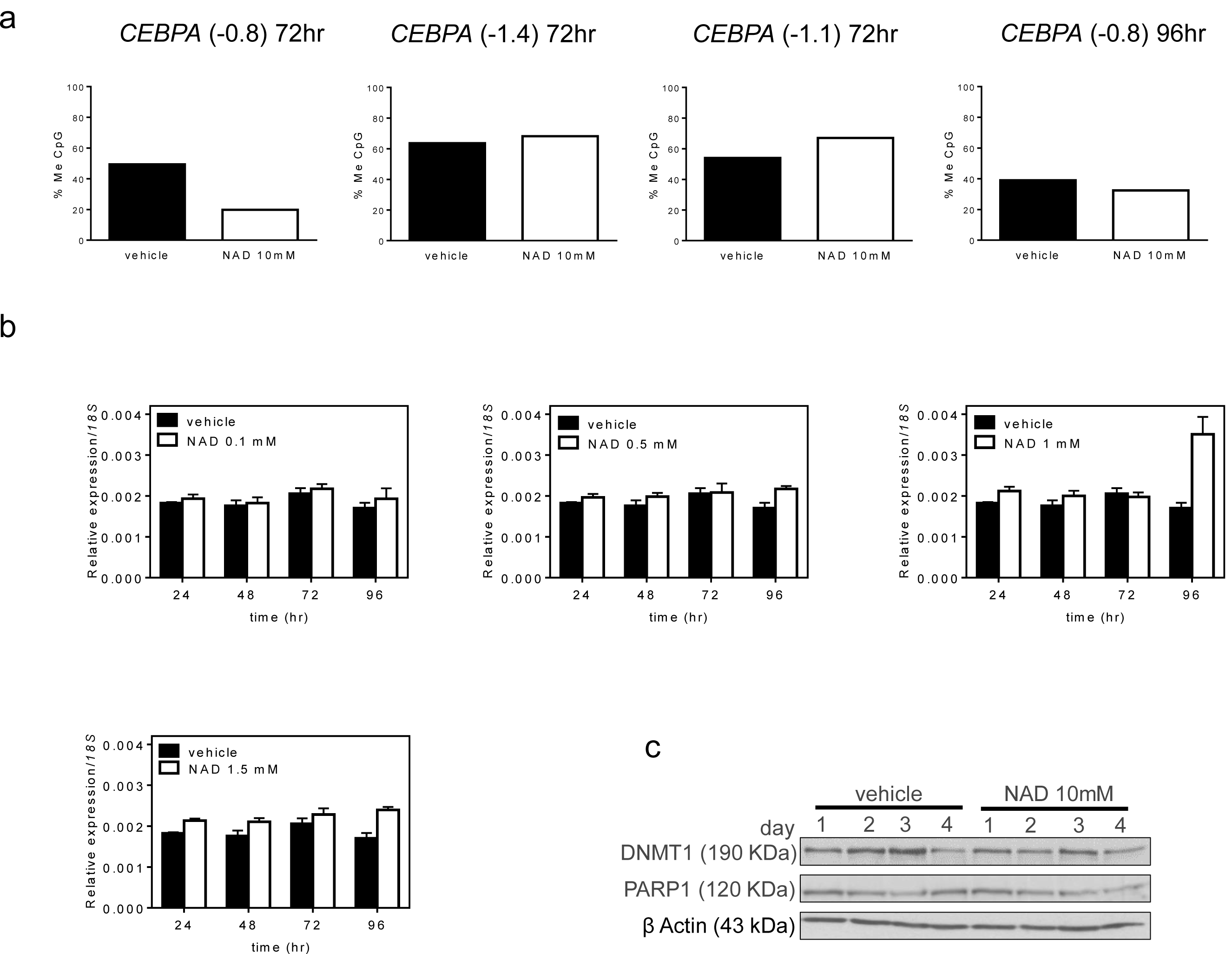
DNA methylation patterns and expression profile of *CEBPA* upon NAD or vehicle treatment. (**A**) Histograms representing the percentages of methylated CpGs (% Me CpGs) across *CEBPA* promoter 72hrs or 96hrs after treatment with NAD (10mM), calculated by Quma software. (**B**) expression levels of *CEBPA* in K562 treated for four days with either vehicle (water) or different concentration of NAD (0.1, 0.5, 1, 1.5 mM). (**C**) PARP1 and DNMT1 protein levels in K562 upon NAD treatment were monitored by western blot analysis.

**Supplementary figure 3.**
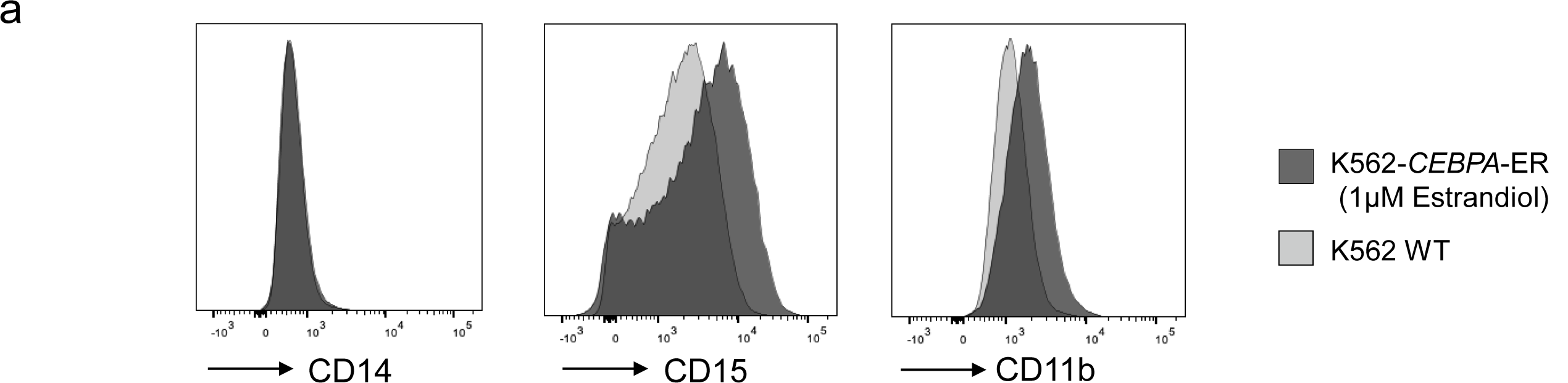
Flow cytometry analysis of CD14, CD15 and CD11b expression, in K562 wild type and K562-*CEBPA*-ER differentiated cells.

