## Supplementary Information for "NAD modulates DNA methylation and cell differentiation"

**Supplementary material**

**Table 1** qRT-PCR primer sets

| Primers | CEBPA |
| --- | --- |
| Forward | 5’-TCGGTGGACAAGAACAG-3’ |
| Reverse | 5’-GCAGGCGGTCATTG-3’ |
| Probe | 5’-ACAAGGCCAAGCAGCGC-3’ |
|  | PARP1 |
| Forward | 5’-AAGATGATCTTTGATGTGGAAAGTA-3’ |
| Reverse | 5’-TGCCCTTGGGGAAGCTGAGCAAA-3’ |
| Probe | 5’-GAAGAAAGCCATGGTGGAGT-3’ |

DNMT1(ThermoFisher cat. No. hs00945875_m1); GAPDH (ThermoFisher cat. No. 4310884E); 18S rRNA (ThermoFisher cat.No. 4310893E).

**Table 2** Bisulfite sequencing primer sets

| Region | Forward Primer | Reverse Primer |
| --- | --- | --- |
| -0.8 (-557; -857) | 5’-CAGCTCCGCTAGTCTGGGGGGCC-3’ | 5’-CACAGGGGTAGCCTGGAGATCAGA-3’ |
| -1.1 (-895; -1.122) | 5’-CACTCAAGGGGCCCCAGG-3’ | 5’-CCAGAGTTAAGTTTGTCTCC-3’ |
| -1.4 (-1.120; -1.473) | 5’-GGTGTTTTTAGCTGTGCCCCCT-3’ | 5’-TCAAGGGGCCCCAGGGCCT-3’ |

**Table 3** ChIP-qPCR primer sets

| Region | Forward Primer | Reverse Primer |
| --- | --- | --- |
| -0.8 (-557; -857) | 5’-CAGCTCCGCTAGTCTGGGGGGCC-3’ | 5’-CACAGGGGTAGCCTGGAGATCAGA-3’ |
| -1.4 (-1.120; -1.473) | 5’-GGTGTTTTTAGCTGTGCCCCCT-3’ | 5’-TCAAGGGGCCCCAGGGCCT-3’ |
